# The E3 ubiquitin-protein ligase MDM2 is a novel interactor of the von Hippel-Lindau tumor suppressor

**DOI:** 10.1101/2020.04.28.066084

**Authors:** Antonella Falconieri, Giovanni Minervini, Raissa Bortolotto, Damiano Piovesan, Raffaele Lopreiato, Geppo Sartori, Maria Pennuto, Silvio C.E. Tosatto

## Abstract

Mutations of the von Hippel-Lindau (pVHL) tumor suppressor are causative of a familiar predisposition to develop different types of cancer. pVHL is mainly known for its role in regulating hypoxia-inducible factor 1-α (HIF-1α) degradation, thus modulating the hypoxia response. There are different pVHL isoforms, including pVHL30 and pVHL19. However, little is known about isoform-specific functions and protein-protein interactions. Integrating *in silico* predictions with *in vitro* and in *vivo* assays, we describe a novel interaction between pVHL and mouse double minute 2 homolog (MDM2). Importantly, we found that pVHL30, and not pVHL19, forms a complex with MDM2, and that the N-terminal acidic tail of pVHL30 is required for its association with MDM2. Further, we demonstrate that an intrinsically disordered region upstream of the tetramerization domain of MDM2 is responsible for its isoform-specific association with pVHL30. This region is highly conserved in higher mammals, including primates, similarly to what has been already proposed for the N-terminal tail of pVHL30. Finally, we show that overexpression of pVHL30 and MDM2 together reduces cell proliferation, suggesting a synergistic effect of these E3 ubiquitin ligases. Collectively, our data support the idea that pVHL30 plays a role in MDM2 regulation, suggesting a wider interplay among hypoxia sensing and cell cycle regulation.

## Introduction

Oxygen availability regulates a plethora of physiological processes, such as cell metabolism, mitochondrial ATP production, embryo development and inflammation, and it plays a prominent role in the pathophysiology of cancer ^1–3^. Hypoxia response in cells is carried out by the hypoxia-inducible factor (HIF) complex in a tightly oxygen-dependent fashion ^4,5^. HIF is a heterodimeric complex formed by an α-subunit, whose stability is finely regulated by the intra-cellular oxygen concentration, and a constitutively expressed nuclear β-subunit ^6,7^. As a transcription factor, HIF is a master activator of many genes that allow the cells to cope with low oxygen growth conditions by enhancing glycolytic flux while inhibiting mitochondrial respiration ^8^. In physiological conditions, the levels of HIF-1α are regulated by the von Hippel–Lindau (pVHL) tumor suppressor, which continuously targets HIF-1α for proteasomal degradation. As a substrate recognition particle of the VCB multiprotein complex (together with Elongin B, Elongin C, the E3-ubiquitin ligase Cullin-2 and RBX1 ^9^), pVHL recognizes two proline residues of HIF-1α^10^ upon hydroxylation by prolyl hydroxylase domain containing proteins (PHD) ^11^, whose expression in turn is closely regulated by HIF-1α itself^12^. In hypoxia, the activity of PHDs is inhibited, resulting in HIF-1α stabilization and activation of the hypoxia-response gene expression program. Together, pVHL, HIF-1α and PHD form the cellular oxygen-sensing system ^4,11,13,14^.

Deregulation of HIF-1α/pVHL promotes sustained angiogenesis ^15^ and is causative of von Hippel-Lindau syndrome ^16–18^, a familiar predisposition to develop different types of cancer, such as pheochromocytoma, paraganglioma, retinal- and cerebellar-hemangioblastoma and renal cell carcinoma (RCC)^19^. In humans, pVHL is coded by the *VHL* gene located on chromosome 3^20^. Three biologically active pVHL isoforms have been described so far, namely pVHL30, pVHL19, and pVHL172. Both pVHL19 and pVHL30 act as tumor suppressors and have redundant function to promote HIF-1α degradation, yet they have a different sub-cellular localization ^21^ and isoform-specific functions ^9,22^. Conversely, rather than playing a role as tumor suppressor, pVHL172 contributes to renal carcinoma by up-regulating a subset of pro-tumorigenic genes ^23^, i.e. *Tgfb1*, *Mmp1*, and *Mmp13*. pVHL30 was proposed to specifically interact with p14ARF ^22^, suggesting a further connection between oxygen sensing and apoptosis. Although these observations suggest that the different pVHL isoforms have redundant as well as isoform-specific functions in the cells, little is known about pVHL isoform-specific protein-protein structural and functional interactions. Here, we report a novel direct interaction between mouse double minute 2 homolog (MDM2) and pVHL30. By undertaking a complementary approach, spanning from *in silico* analyses to yeast two-hybrid and mammalian cell assays, we defined the molecular details driving this novel interaction. Finally, we showed that a short intrinsically disordered segment of MDM2 is responsible for its association with pVHL in an isoform-specific fashion restricted to pVHL30.

## Results

### *In silico* protein-protein interacting network analysis identifies MDM2 as a novel interactor of pVHL

We previously documented a binary interaction between pVHL30 and p14ARF sustained by the acidic N-terminal tail of pVHL30, suggesting a functional asymmetry among pVHL isoforms ^22^. p14ARF tumor suppressor is well known for its role in regulating p53 stability^24^, sequestering MDM2 into a subnuclear compartment ^25^, while inhibiting p53 degradation ^24^. p53 is also stabilized through pVHL ^26^. In Zebrafish MDM2 degradation was regulated upon pVHL interaction with programmed cell death 5 protein (PDCD5), a negative regulator of MDM2. This evidence suggests a direct functional effect of pVHL loss of function on the p53/MDM2 pathway. To address this question, we used STRING to generate an interaction network centered on pVHL, HIF-1α, p14ARF, p53, and MDM2 ^27^. The resulting network was composed of twenty nodes connected by 52 edges, each of them representing an experimentally validated interaction (Figure 1). The network is significantly more connected than expected by chance (p-value < 1.0e^−16^), implying that these proteins are functionally interconnected. We asked whether non-trivial functional subnetworks can be linked to specific cell functions. We identified three distinct clusters formed by proteins linked to apoptosis, protein ubiquitination and hypoxia response. The first cluster was formed by six proteins, including p53, MDM2, and p14ARF. Gene ontology terms enrichment predicts this cluster to be mainly localized in the nucleus and participating in the regulation of transcription in response to hypoxia and cellular stress. The second cluster was composed of seven proteins, including pVHL, and, as expected, it was linked to protein degradation. The last cluster was the most heterogeneous and included proteins associated with both hypoxia- and stress-response, e.g. nutrient deprivation and viral aggression. Of note, we found a number of experimentally validated interactions connecting all of these three clusters, suggesting that the hypoxia response is functionally interconnected with cell cycle regulation. Manually curated enrichment of protein-protein interactions across network nodes further extended the connection between clusters I and II. Considering the concept of “interactors of interactors”, a well-established property of protein-protein interaction networks ^28,29^ and the already validated physical interaction between pVHL, p53, and p14ARF, our *in silico* analysis predicts that MDM2 is a novel direct interactor of pVHL.

**Figure 1.**
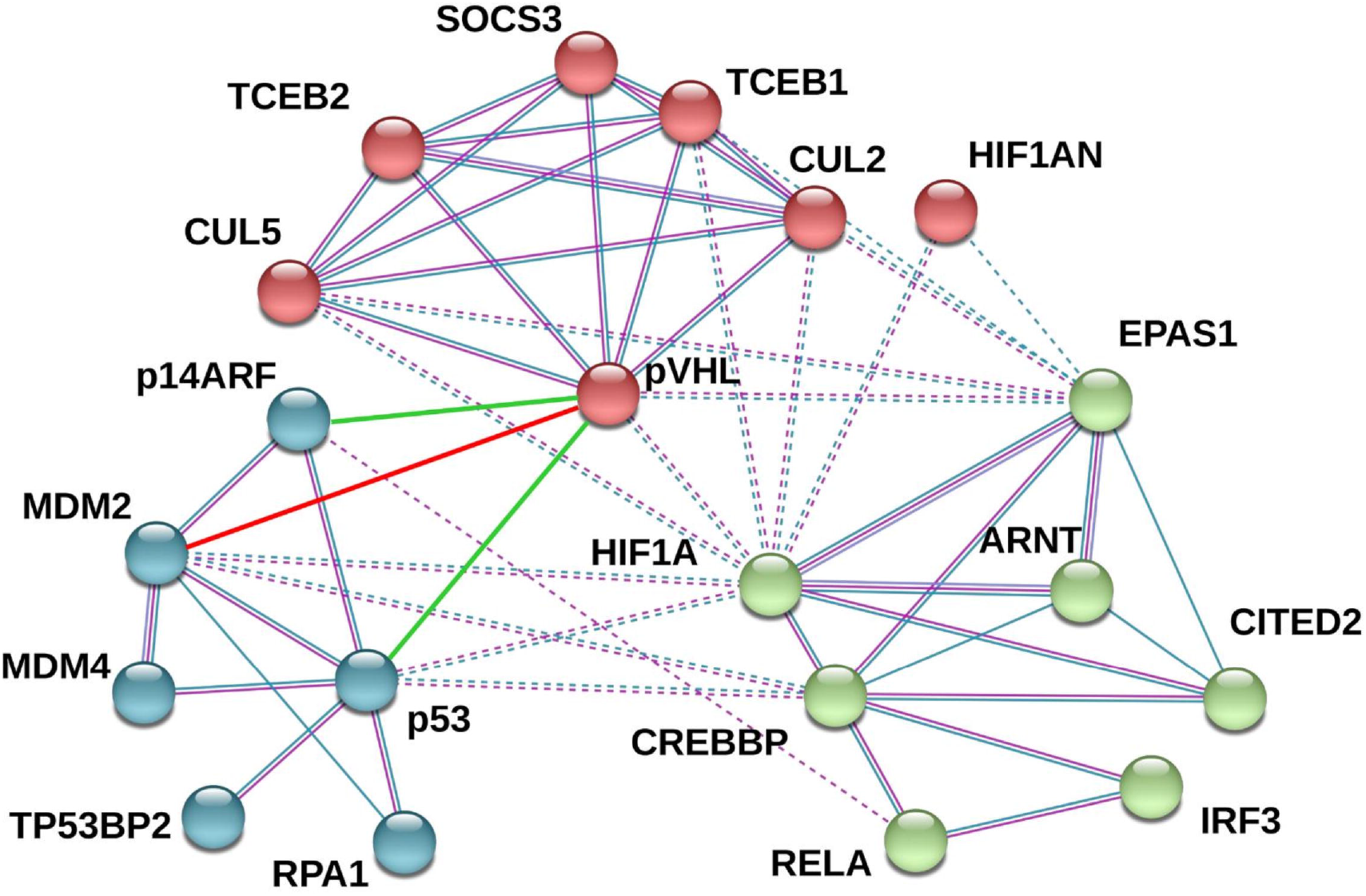
Network of interacting proteins. Connections between nodes represent experimental evidence for interaction, while colored bubbles are for proteins forming the network. A green edge represents the interaction between pVHL30 and p14ARF (CDKN2A), currently not yet reported in STRING, while a red line is used for the novel association of pVHL30 with MDM2 reported here.

### pVHL30 specifically interacts with MDM2

To validate the *in silico*-resolved pVHL association with MDM2, we used yeast two-hybrid (Y2H) assays. We tested whether pVHL binds to MDM2 in an isoform-specific fashion. To address this question, we generated vectors to express human MDM2 fused with Gal4-binding domain (Gal4-BD) as bait, and human pVHL30 and pVHL19 fused with Gal4-activation domain (Gal4-AD) as prey. Protein expression in yeast cells was verified by Western blotting (Figure S1). Transient expression of either pVHL30, pVHL19, or MDM2 alone did not result in yeast growth in selective medium. On the other hand, co-expression of pVHL30 and MDM2 together resulted in an increase in yeast cell growth in selective medium, thereby demonstrating the ability of MDM2 and pVHL30 to associate with each other in yeast cells (Figure 2A-B and Figure S2). Although Y2H data cannot be considered quantitative *per se*, these results suggest a weak interaction, as the yeast growth rate was less sustained compared to the positive control. Importantly, co-expression of pVHL19 and MDM2 did not result in cell growth in selective medium, indicating that pVHL19 is unable to associate with MDM2 under these experimental conditions.

**Figure 2.**
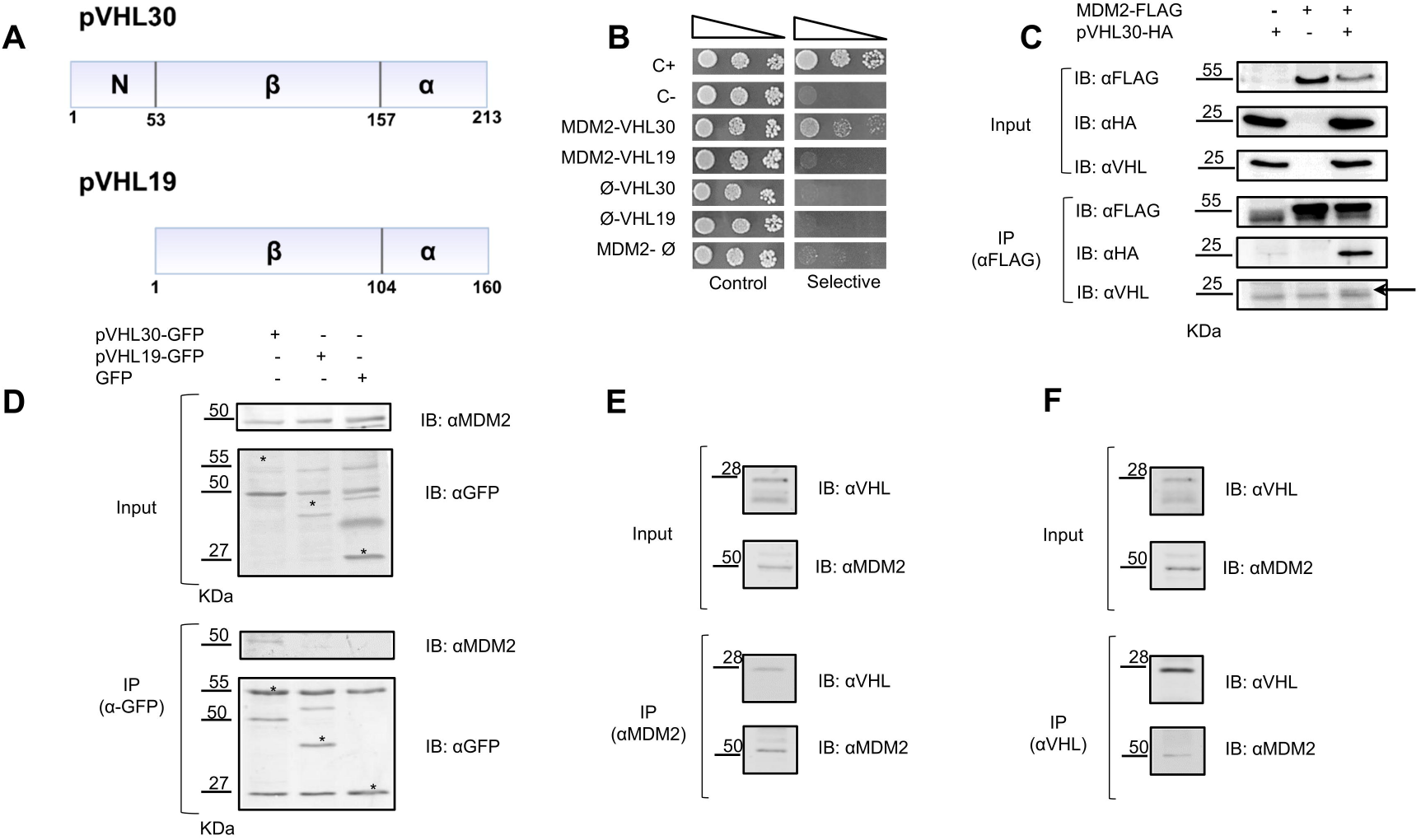
pVHL interacts with MDM2 in an isoform-specific manner. A) Schematic representation of pVHL isoform domain composition. The two isoforms differ for the presence of an intrinsically disordered N-terminus. B) Y2H assay showing that only pVHL30 associates with MDM2. C+, positive control. C−, negative control. C) Immunoprecipitation (IP) of total protein extracts from HEK293T cells expressing Flag-MDM2 and HA-VHL30 with an anti-Flag antibody and immunostained with anti-Flag antibody, and anti-pVHL/anti-HA antibodies, as indicated. D) IP of total protein extracts from MN1 cells expressing pVHL30-GFP, pVHL19-GFP, and soluble GFP, as indicated, and processed for pull down with anti-GFP antibody revealed endogenous MDM2 pull down only in the presence of pVHL30-GFP. Endogenous MDM2 was detected with a specific antibody. E-F) IP of total protein extracts from HEK293T cells with either an anti-MDM2 antibody (E) or anti-pVHL antibody (F) showed reciprocal pull down of endogenous proteins.

Next, we asked whether pVHL binds to MDM2 in an isoform-specific manner in mammalian cells. To test this hypothesis, we used human embryo kidney (HEK) 293T cancer cells transiently transfected with vectors expressing either HA-tagged pVHL30 or pVHL19 with Flag-tagged MDM2. Using immunoprecipitation and Western blotting analysis, we found that pVHL30 forms a complex with MDM2 (Figure 2C and Figure S3). However, using this approach we could not establish whether pVHL19 forms a complex with MDM2 in mammalian cells (Figure S4). To determine whether pVHL30 binds endogenous MDM2 in an isoform-specific fashion, we generated vectors to express green fluorescent protein (GFP)-tagged pVHL30 and pVHL19. Overexpression of pVHL30-GFP, and not pVHL19-GFP, in motor neuron-neuroblastoma hybrid MN-1 cells resulted in the pull down of endogenous MDM2 (Figure 2D and Figure S5). Finally, we asked whether endogenous pVHL forms a complex with endogenous MDM2 (Figure 2E-F and Figure S6). Immunoprecipitation of endogenous pVHL resulted in the pull down of endogenous MDM2. Notably, reciprocal immunoprecipitation of MDM2 specifically pulled down the pVHL isoform with the higher molecular weight, which corresponds to pVHL30. These observations indicate that pVHL30 specifically forms a complex with MDM2 in yeast as well as human- and mouse-derived mammalian cancer cells. Moreover, our results suggest a functional role of the pVHL30 N-terminal tail in driving the interaction with MDM2.

### MDM2 C-terminal domain binds to pVHL30

Next, we sought to determine which region of MDM2 specifically binds pVHL30. We generated vectors expressing MDM2 N-terminus (1-150), central acidic domain (151-350), and C-terminus (351-491) (Figure 3A). Expression of truncated MDM2 proteins in yeast was confirmed by Western blotting (Figure S1). Yeast cells co-expressing pVHL30 together with the N-terminus or central acidic domain of MDM2 did not grow in selective medium, indicating no binding (Figure 3B and Figures S7). Rather, yeast cells co-expressing pVHL30 and the MDM2 C-terminus grew in selective medium. These results indicate that the last 140 C-terminal residues of MDM2 (MDM2-CT, aa 351-491) are key for the pVHL30/MDM2 interaction. Based on these observations we further characterized the C-terminal domain of MDM2. Analysis of the MDM2 primary structure with FELLS revealed hallmarks of an intrinsically disordered region spanning residues 329 to 433, terminating before the C3HC4-type zinc finger domain (RING, aa 434-491), which mediates MDM2 protein-protein and protein-RNA interactions ^30–32^ (Figure 3C). By a Y2H assay, cells co-expressing both RING domain and pVHL30 did not grow in selective medium, indicating that the MDM2 RING domain does not associate with pVHL30 (Figures 3D-E and Figure S8). Yeast cells co-expressing pVHL30 and the MDM2 351-433 fragment also failed to grow in selective medium. Rather, the interaction with pVHL30 was restored by slightly elongating the C-terminal end of this fragment to include the first part of the RING domain, as indicated by the survival of yeast cells co-expressing pVHL30 and MDM2 351-451, despite the overall disruption of the RING domain structure. Collectively, these findings suggest that at least a portion of the RING domain mediates and/or modulates MDM2-pVHL30 association.

**Figure 3.**
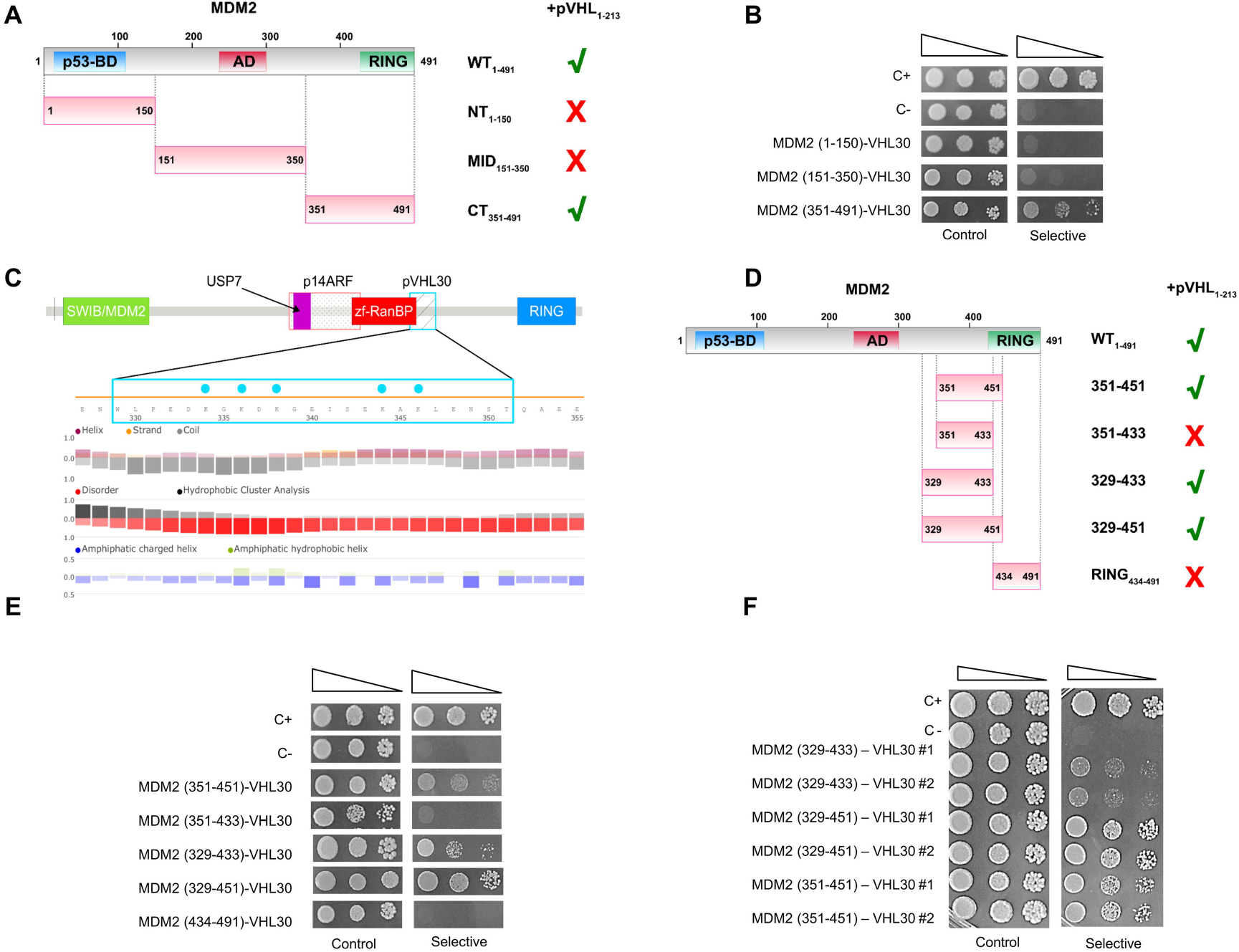
The intrinsically disordered portion of MDM2 is required for pVHL30 binding. A) Schematic representation of MDM2 domain composition. B) Y2H assays revealed that the C-terminal portion of MDM2 binds pVHL30. C) Zoomed view and FELLS representation of the binding motif driving pVHL30/MDM2 association. Blue dots represent positively charged residues. The identified binding region partially overlaps the pseudo zinc-finger zf-RanBP. D) Schematic representation of the C-terminal region MDM2 fragments used in panels (E-F). E-F) Y2H assays revealed that the interaction of pVHL30 with MDM2 requires either the 329-350 or 434-451 regions of MDM2, yet the presence of the entire region (329-451) enhanced yeast growth.

We thus wondered whether the MDM2 disordered region (aa 329-433) mediates the interaction with pVHL30, possibly by an additional and/or cooperative mechanism within the MDM2 C-terminus. Considering that the pVHL30 N-terminal tail, which is specifically involved in this interaction, is mostly acidic and characterized by the presence of eight GxEEx repetitions ^22^, we investigated whether clusters of positively charged residues within the MDM2 disordered region account for pVHL30 binding. *In silico* characterization with FELLS identified a region spanning residues 329-351 as a good candidate for pVHL30 binding (Figure 3C). This region is predicted as fully disordered, with a typical alternation of charged residues. Of note, the position of a stretch of five lysine residues suggests these to form a positively charged cluster, which we hypothesize to form electrostatic interactions with the acidic pVHL30 N-terminus. In Y2H assay, co-expression of pVHL30 and the MDM2 329-433 fragment resulted in cell growth in selective medium, indicating that residues 329-433 are the minimal MDM2 sequence required for interaction with pVHL30 (Figures 3D-E and Figure S8). These data highlight the relevance of MDM2 residues 329-350 for binding, as interaction with pVHL30 is completely lost upon their removal. On the other hand, the stretch 329-350 was not sufficient for MDM2-pVHL30 interaction, as the MDM2 151-350 fragment was unable to bind pVHL30 (Figure 3B). Interestingly, co-expressing pVHL30 with a longer MDM2 fragment (aa 329-451), which includes a portion of the RING domain implicated in the interaction (434-451), enhanced the growth rate of yeast cells in selective medium (Figure 3E), supporting the idea that these two flanking sequences cooperatively improve the binding stability. To confirm this hypothesis, we further compared the growth of yeast cells co-expressing pVHL30 with MDM2 329-433, 329-451, and 351-451 fragments (Figure 3F). Yeast cells grew more in presence of MDM2 329-451, pointing to a role of the flanking sequences to the binding stability/regulation.

### The pVHL30 N-terminus and β-domain are required for interaction with MDM2

pVHL has multiple binding surfaces ^33^, each responsible for specific protein-protein interactions ^34^. To identify the pVHL30 surface required for MDM2 binding, we generated yeast vectors expressing the pVHL N-terminus (aa 1-53), β-region (aa 54-157), and C-terminal α-domain (aa 158-213), and we performed Y2H assays. Expression of these pVHL30 truncated proteins in yeast was confirmed by Western blotting (Figure S1). The N-terminus and β- and α-regions did not associate with MDM2 full length (Figure S9) and MDM2 329-433 fragment (Figure 4A-B and Figure S10). These findings show that the pVHL30 N-terminal tail, although necessary (Figure 2B), is not sufficient for MDM2 binding. Rather, our data suggest that pVHL30/MDM2 interaction is cooperatively driven by concurrent participation of the intrinsically disordered N-terminal tail and at least another pVHL30 domain. Consistent with this hypothesis, a Y2H assay showed that yeast cells expressing the pVHL30-N-β fragment (1-157) containing both the acid tail and ß-domain together with MDM2 329-433 fragment grew in selective medium, indicating that these portions of pVHL30 and MDM2 are critical binding surfaces required for such interaction (Figure 4C-D). Taken together, this experimental evidence supports the idea that pVHL30/MDM2 interaction occurs through the MDM2 C-terminal region (aa 329-491) combining the binding of multiple interaction motifs within the 350-433 region to the pVHL-β domain. We propose that the pVHL30/MDM2 interaction relies on two factors, on one side the electrostatic interactions of the positively charged MDM2 residues (localized in the 329-350 stretch) with the negatively charged pVHL-N-terminus, and on the other side the positive contribution of (or part of) the RING domain.

**Figure 4.**
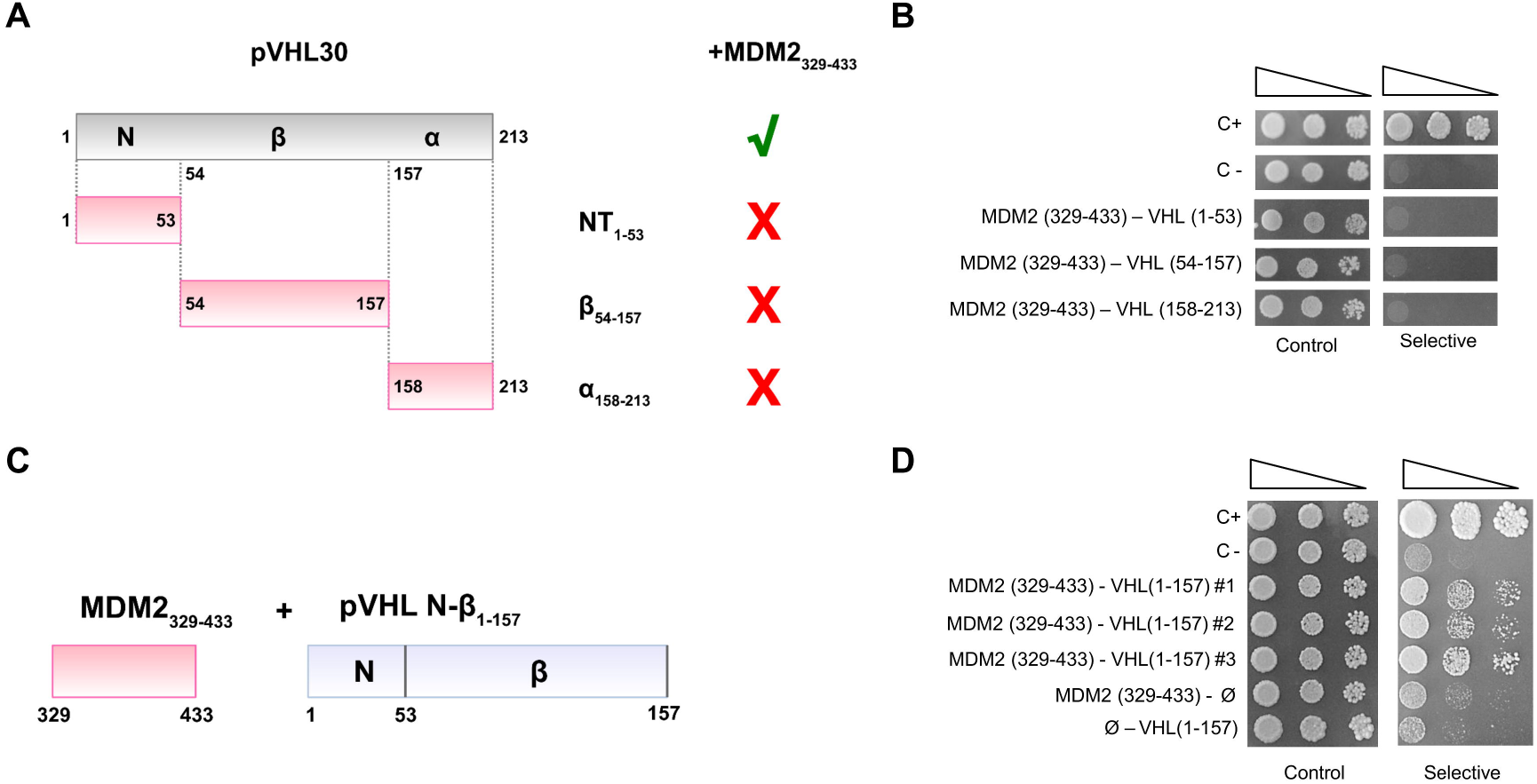
The N-terminus and β-domain of pVHL30 are required for MDM2 binding. A) Schematic representation of pVHL30 domain composition. B) Y2H assays revealed that none of pVHL30 domains alone bind MDM2 fragment 329-433. C) Schematic representation of the MDM2 and pVHL30 fragments used in panel (D). D) Y2H assays showed the interaction of pVHL30 1-157 region with MDM2 329-433 region.

### *Ab initio* structure prediction and intrinsically disorder flavors characterization of MDM2 fragments

Limited structural information is available about the MDM2 region mediating the association with pVHL30 as no crystal structure is currently available for this region. Since intrinsically disordered regions, albeit lacking a fixed tertiary structure, are often associated with protein cellular function(s) depending on chemical property intrinsic to their primary structure ^35–37^, we investigated whether structural elements in MDM2 explain the different binding behaviors of MDM2 regions. To this aim, we performed *ab initio* structure prediction and disorder flavors characterization of the different MDM2 fragments tested in yeast. All predictions consistently showed that the MDM2 fragments spanning 351-451, 351-433, 329-433, and 329-451 are mostly disordered, with relevant differences in secondary structure content (Figure 5A and Figure S11). The MDM2 351-451 fragment, containing the minimal sequences required for association with pVHL30, is characterized by a relatively high content of secondary structure. A small β-hairpin followed by a variable number of alpha bulges seems to be shared among different predictions. The β-hairpin is lost in predictions of MDM2 329-433 (which binds pVHL30), while alpha bulges are conserved. MDM2 329-451 is characterized by unfolded-like conformations albeit presenting few conserved secondary structure elements. On the other hand, the shorter MDM2 351-433 fragment, which lacks both flanking regions promoting MDM2-pVHL30 binding, is predicted as the most compact. Average secondary structure content, calculated upon 5,000 3D structure predictions for each fragment, indicated that alpha bulges are conserved in both interacting and not interacting MDM2 fragments. These bulges mostly localize in the stretch 351-433, with flaking regions more frequently predicted as coil (Figure 5B). Detailed characterization of residue-residue interactions for each model performed with RING ^38^ shows that the MDM2 351-433 (unable to bind pVHL30) is characterized by the highest hydrogen bonds, Van der Waals and ionic interactions (Figure 5C). This finding supports the concept that this region, when isolated from flanking regions, acquires a more globular-like conformation. Conversely, on average the other fragments has comparable hydrogen bonds paired with a lower number of Van der Waals and ionic interactions, thus being relatively rich in secondary structure and more prone to adopt extended conformations. Collectively, all of these findings suggest that shortening MDM2 329-451, by removing both N- and C-flanking regions, stabilizes the polypeptide, thus lowering the intrinsically disordered behavior and inducing a conformational switch to order. Of note, multiple sequence alignment shows that the positively charged MDM2 329-351 region is maintained throughout evolution and is mostly conserved in Artiodactyla and Primates (Figure S12). As the number of acidic repeats forming pVHL30 N-terminal tail increases from rodents to primates ^22^, data suggest that the isoform-dependent MDM2/pVHL30 association could have specifically evolved in higher mammals. We further observed that the cluster of positive charges in region 329-351 appears counterbalanced by an opposed concentration of negative charges in region 410-433 (Figure S12), supporting the hypothesis that the two flanking sequences interact with the pVHL30 acidic N-terminus and β-domain, respectively, as pVHL β-domain presents a number of conserved positive residues mediating multiple protein-protein interactions ^10,33,28^. Notably, MDM2 residues spanning 351-433 fragment includes six ATM (ataxia-telangiectasia mutated kinase) phosphorylation sites (i.e., S386, S395, S407, T419 S425, S429, Figure S12), strongly suggesting that post-translational modifications may operate as regulator of MDM2/pVHL30 association, e.g. introducing electrostatic repulsion against the pVHL30 acidic tail upon phosphorylation.

**Figure 5.**
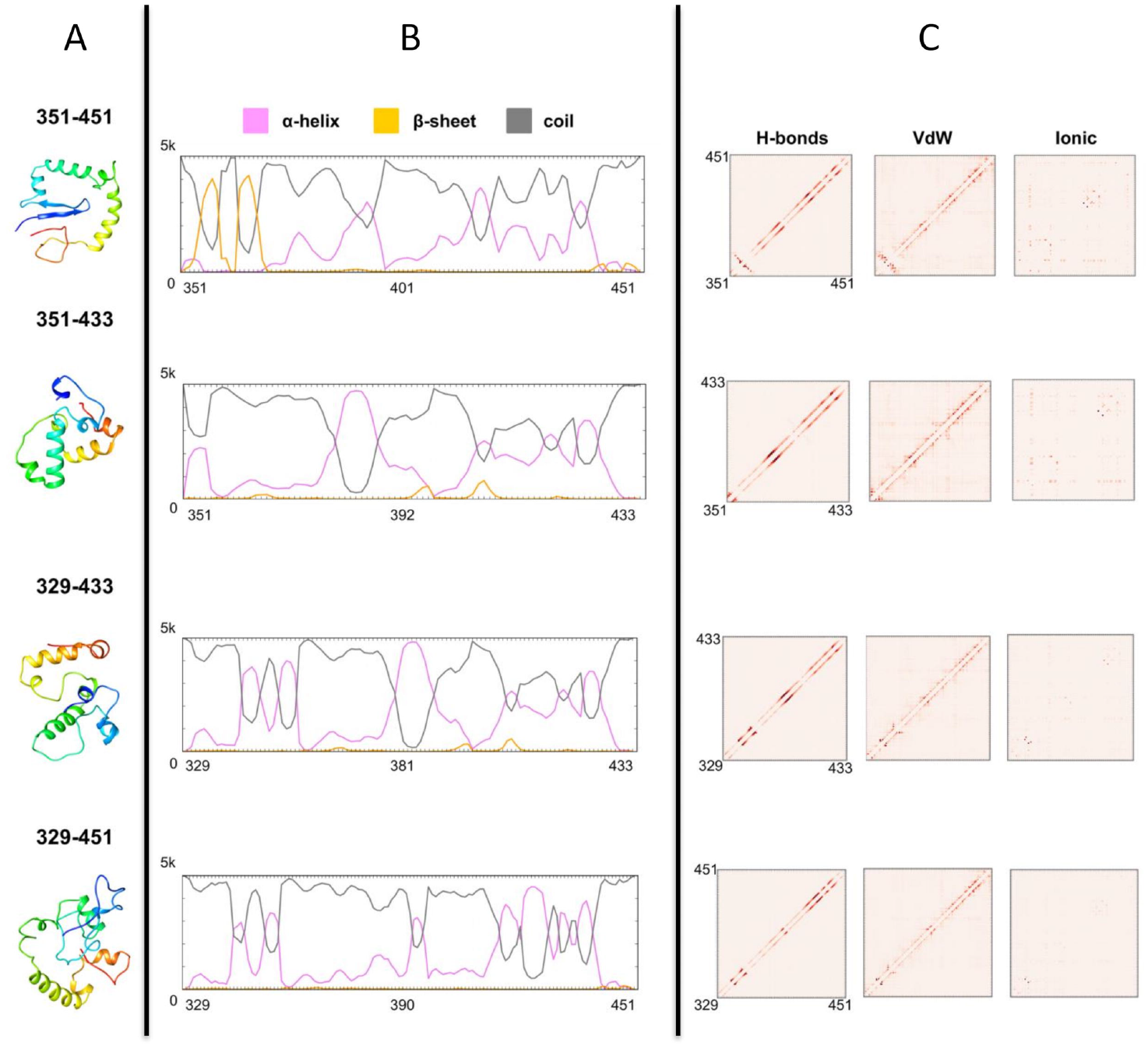
Characterization *in silico* of structural features of MDM2 fragments. Ab initio structure predictions (A), secondary structure content (B) and conserved intra-chains interactions (C) are presented for each fragment. The most probable 3D structures are presented as cartoon and colored rainbow with blue and red marking the N- and C-terminus respectively. Secondary structure content is calculated analyzing 5,000 predictions is presented as plot. Intra-chain interactions describing network of interacting residues and type of interactions are presented as boxplots.

### pVHL30 and MDM2 reduce cell proliferation

MDM2 is a E3 ubiquitin-protein ligase^39^ that mediates p53 degradation^40^. MDM2 physiological activity establishes a regulative feedback loop that maintains optimal levels of p53^40–42^, thus modulating activation of apoptosis upon cellular stress^43^. To gain insights into the biological relevance of pVHL30/MDM2 interaction, we performed a cell proliferation assay in MN1 cells transiently transfected with vectors expressing HA-tagged pVHL30 and Flag-tagged MDM2 (Figure 6). We found that overexpression of either pVHL30 or MDM2 alone did not modify cell proliferation. Rather, co-expression of both E3 ubiquitin-protein ligases significantly reduced cell proliferation by 19%. These data suggest a cooperative and synergistic effect of these E3 ubiquitin-protein ligases in the regulation of cell proliferation.

**Figure 6.**
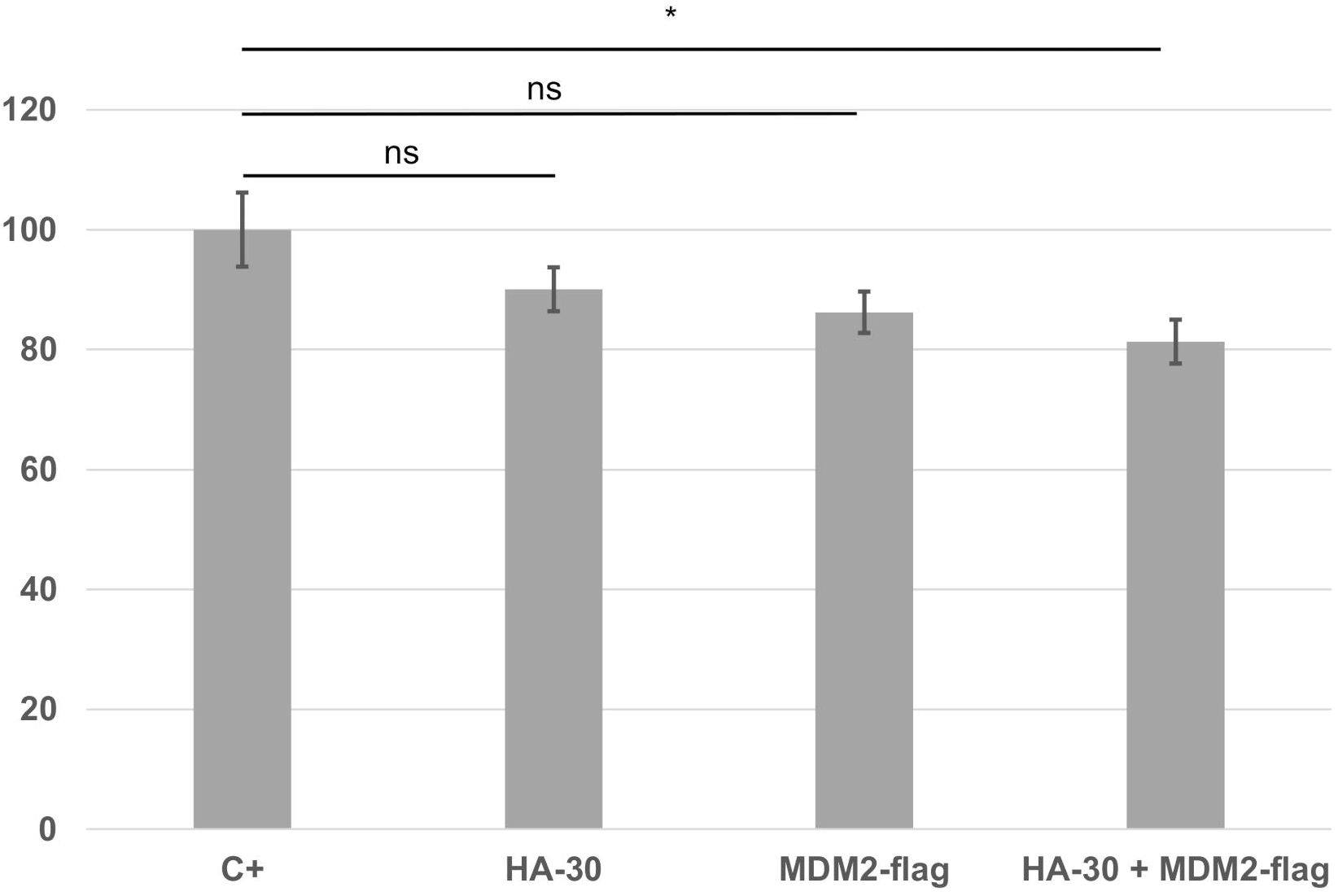
Gain of pVHL30 and MDM2 function reduces cancer cell proliferation. MTT assay in MN1 cells transiently transfected with vectors expressing pVHL30 and MDM2, as indicated, revealed that overexpression of both E3 ubiquitin ligases together significantly reduces cell proliferation. Graphs, mean +/− sem (n = 12), ANOVA followed by Bonferroni post-hoc, * p < 0.05.

In summary, our results show an isoform-specific pVHL protein-protein structural and functional interaction. Together with our previous finding of pVHL30-p14ARF specific interaction ^9,22^, these observations support the idea that pVHL30 and the other isoforms have non-redundant functions in the cells.

## Discussion

Functional intersections between oxygen sensing, apoptosis and cell cycle regulation are well described in the literature ^26,28,44,45^, although their functional inter-connection and mutual regulation are still debated. In particular, under prolonged hypoxia, p53 has been reported to accumulate in the cell, yielding a direct repression of HIF-1α transcriptional activity ^46^. A ternary complex between p53, HIF-1α, and MDM2 was proposed to promote pVHL-independent degradation of HIF-1α ^44^. pVHL has been shown to directly associate with p53, thereby inducing both suppression of MDM2-mediated ubiquitination and nuclear export of p53 ^26^. Degradation of p53 is also regulated by p14ARF tumor suppressor ^24^, which inhibits the MDM2 E3 ubiquitin-protein ligase activity, leading to p53 accumulation and apoptosis ^47^. Here, we describe a novel direct interaction between pVHL and MDM2. This association is pVHL30-isoform-specific and was observed in yeast and mammalian cells. Importantly, we provide evidence that this interaction occurs through the intrinsically disordered N-terminal tail of pVHL30, which indeed is not present in pVHL19. Notably, this acidic tail is necessary, but not sufficient and requires the pVHL β domain. This domain is known to engage a number of different protein-protein associations ^34^, and consistent with this idea it harbors the HIF-1α binding site. HIF-1α associates with MDM2 ^44^, an interaction thought to play a role in modulating hypoxia-induced p53 activation. Our data, while pointing out a novel interaction between MDM2 and pVHL30, suggest that the two pVHL isoforms evolved to absolve different functions. We speculate that transient macromolecular complexes composed of proteins participating in the regulation of both hypoxia sensing and cell cycle regulation may constantly be formed and destroyed in different cell compartments. Ideally, under mild hypoxia conditions the increased amount of free pVHL may trigger a pVHL30 functional relocation and MDM2 association. Of note, the MDM2 region that we found to participate in this association is immediately downstream of two overlapping binding sites for p14ARF (aa 210-304) and USP7 (aa 242-331), a deubiquitinating enzyme involved in p53 stabilization ^48^. Within the MDM2 351-433 region, multiple phosphorylation sites for the ATM kinase are found to be relevant for activation of the DNA damage response ^49^. Our data show that the pVHL30/MDM2 interaction is sustained by at least two linear interacting peptides (LIPs) ^50^ present on the MDM2 *ab initio* models and putatively modulated by either indirect (e.g. variation of oxygen concentration) and/or direct (e.g. site-specific phosphorylation) mechanisms. In the context of transient complexes, the observation that a MDM2 motif driving its association with pVHL30 is immediately adjacent to other binding sites, and it can also be modified by PTM is intriguing. It has been proposed that the interaction of pVHL30 with p14ARF induces its nuclear-to-nucleolar translocation, with concurrent dissociation of pVHL30 from the pVHL-elongin C-elongin B (VCB) complex ^9^. A similar mechanism also triggers p14ARF-dependent functional inhibition of MDM2 ^47^. VCB-unbound pVHL30 can interact and stabilize p53 through its α-domain, by suppressing Mdm2-mediated ubiquitination as well as nuclear export ^26,34^. Here, we demonstrated that both the N-terminal tail and β-domain are necessary for interaction with MDM2. Such kind of association involves different regions of the protein and is common among linear motif-mediated protein-protein interactions ^51,52^. When bound to MDM2, pVHL30 may conceal ATM-specific phosphorylation sites yielding a further regulative layer for MDM2 activity. Indeed, it has been already demonstrated that ATM induces p53 accumulation and activation by regulating MDM2 oligomerization ^49^, while pVHL associates with ATM impairing MDM2-mediated ubiquitination and p53 nuclear export ^26^. Acting as a protein-hub, pVHL30 may concurrently form alternative ternary complexes under different conditions. The presence of an USP7 binding site in the same MDM2 region suggests that deubiquitination of several target proteins, such as MDM2 and p53, promotes their stabilization ^48,53^. Consistent with this idea, other proteins can mediate USP7 association with p53 ^54^. In our recent investigation ^22^, we found two putative USP7 binding sites on the pVHL30 N-terminus (residues 35–44). Collectively, we propose that in addition to suggesting a novel HIF-1α independent function, the pVHL30 association with MDM2 may play a novel role in MDM2 stabilization and mutual regulation between oxygen sensing and apoptosis.

## Methods

### Bioinformatics analysis

A network of interacting proteins was built with Cytoscape ^55^ retrieving interaction data from STRING ^27^ (medium confidence 0.400), imposing no more than 5 interactors for the first shell and 10 for second shell. To reduce the number of false positive, text mining derived data were manually removed. The final network is composed of 20 nodes connected by 52 edges. The network shows an average node degree of 5.2 and an average local clustering coefficient of 0.721, with expected number of edges of 25. Human MDM2 and pVHL sequences were retrieved from UniProt ^56^ and visualized with Jalview ^57^. Orthologous sequences were retrieved from OMA browser and aligned with a T-Coffee ^58^ (default parameters). Sequence features and secondary structure content were investigated with FELLS ^59^, functional domains mapped on the sequence using InterPro ^60^ and Pfam ^61^. Linear motifs were predicted with ELM ^62^ and intrinsic disorder with MobiDB 3.0 ^63^. Ab initio protein structures prediction was performed with Rosetta 3.8 ^64^ using specific protocol to model intrinsically disordered regions ^65^. For each fragment 5,000 decoys were generated and clustered using Rosetta clustering module. DSSP ^66^ was used to calculate secondary structure content for each 3D structure model, while networks of interacting residues were predicted with RING 2.0 ^38^

### Plasmid constructs

The recombinant plasmids pGADVHL30, pGADVHL-Nt, pGADVHL-β and pcDNA3.1 HA-VHL30 were constructed as previously described ^28^. Genes encoding the human MDM2 and pVHL30 proteins were obtained from commercial plasmid provided by GenScript (GenEZ plasmid OHu28568 and OHu23297) and cDNA transferred into pGBKT7 and pGADT7 plasmids (Clontech) to perform yeast two-hybrid assays (Y2H). PCR was used to amplify cDNAs using primers showed in Table S1 containing 15 nucleotides long 5’ ends addressing precise regions around EcoRI site in the MCS of both pGADT7 and pGBKT7 vectors. PCR obtained were cloned into pGADT7 or pGBKT7 linearized using EcoRI enzyme, following the In-Fusion HD Cloning Kit (Clontech) protocol. All Y2H recombinant plasmids constructed (Table S1) express the proteins in fusion with the Gal4 DNA binding domain (DBD) or the Gal4 DNA activation domain (AD) expressed by pGBKT7 and pGADT7, respectively. Chimeric proteins in yeast cells were revealed using anti HA and cMyc antibodies. GeneScript plasmid OHu28568 was directly used to overexpress MDM2 in mammalian cells, as the cDNA 5’-end is associated with the FLAG epitope. For Co-immunoprecipitation experiments, the pVHL30 and pVHL19 cDNAs sequences were cloned in the BamHI/EcoRI sites of the pcDNA3.1 vector, plus a N-terminal HA epitope sequence. The recombinant plasmids were used to overexpress pVHL30 and pVHL19 as N-terminal HA-tagged proteins. Sanger sequencing was used to verify all cloned sequences.

### Yeast two-hybrid (Y2H) assays

The Matchmaker Gold Two Hybrid System (Clontech) was used to detect direct interaction between MDM2 and pVHL (default conditions). Positive interactions were revealed on multiple conditions by using selective medium lacking histidine and supplemented with 30, 60 or 90 mM 3-AT. Y190 cells were co-transformed with pGBKT7 and pGADT7 empty plasmids as negative control, whereas positive control expresses pGADT7-T (Gal4 AD-SV40 large T-antigen) and pGBKT7-53 (Gal4 DBD-murine p53, fragment 72-390). To exclude erroneous interpretation (i.e. auto-activation), Yeast Y190 cells co-expressing both fusion protein with Gal4 DBD and AD domain alone (and vice-versa) were tested. Expression of the Gal4-fusion proteins in yeast cells has been checked by Western blot analysis (Figure S1). For each transformation three independent clones were serially diluted (10-fold) and spotted either on solid selective medium lacking histidine (30 mM 3-AT and 60 mM 3-AT) or permissive medium, to check cell growth. Yeast strain growth was constantly monitored for 3 to 8 days at 30°C. Total yeast proteins have been obtained by TCA-based solubilization of yeast cells ^67^, followed by standard Western Blot analysis ^68^, using either anti-HA (Abcam, ab16918), or anti-Myc (Abcam, ab127421) antibodies, to reveal the Gal4AD-pVHL or the Gal4BD-MDM2, respectively. Binding of two specific proteins is evaluated by co-expressing the proteins fused with either DNA-binding domain (bait), or the activation domain (prey), of the Gal4 transcription factor. Physical association between bait and prey allows activation of the reporter gene leading to the yeast cell growth on selective medium lacking of histidine.

### Transfection and co-immunoprecipitation from HEK293T cells

Co-immunoprecipitation experiments were performed using human kidney HEK293T cells provided by (Sigma-Aldrich). Cells were plated in a 1 ml well microplate and transfected at 70-80% confluence using Lipofectamine 2000 following the manufacturer protocols (Invitrogen). Each transfection was performed using approximately 10 μg of total DNA (i.e. pcDNA3.1-derived plasmids, empty and/or expressing either HA-pVHL30/pVHL19 or FLAG-MDM2. After 48 hours, transfected cells were washed with phosphate-buffered saline (PBS) and lysed in lysis buffer (20mM HEPES-Na pH7.4, 150mM NaCl, 5 mM CHAPS) supplemented with 1X PIC (Protease Inhibitors Cocktail, Sigma). Cell lysates were centrifuged (10’ at 600 rpm, 4°C) and the resulting post-nuclear supernatant (PNS) quantified by Bradford assay. For co-immunoprecitation, 0.2mg of PNS were incubated for 4 hours at 4°C with 5 μl of protein A magnetic beads (Pierce Thermoscientific) pre-incubated (1 hour at RT) with 2 μg of anti-FLAG antibody (Sigma, F7425). Immune complexes were finally washed 3 times with lysis buffer and eluted by incubating the beads (5’ at 70°C) in 30 μl in 1X NuPAGE LDS sample buffer (Invitrogen) plus 0,1 M DTT. Both PNS and immunoprecipitated samples were separated by SDS-polyacrylamide gel electrophoresis and transferred to nitrocellulose membranes. The membranes were incubated with the anti-HA (Abcam, ab16918), anti-VHL (Santa Cruz, sc-5575) or anti-FLAG (Abcam, ab45766) antibodies to reveal the pVHL30/pVHL19 or MDM2, respectively. Additional Co-IPs were performed overexpressing GFP constructs in MN-1 cells and transfected with Lipofectamine2000 according to manufacturer protocol. Each transfection was performed using 2 μgr of pcDNA3.1-VHL30gfp, pcDNA3.1-VHL19gfp or pcDNA3.1 gfp respectively. All protocol steps as described above. In this case 400 μgr of PNS were incubated with 2 μg of anti-GFP (Abcam, ab1218). Following Western blot, membranes were incubated with anti MDM2 (Sigma, M7815) and anti GFP (Abcam, ab1218). Co-IPs of endogenous proteins was performed using 500 μgr of PNS incubated with 2 μg of anti-MDM2 or 1 μg of anti-VHL (Cell Signaling technology,68547). The following antibodies were used Anti-GFP ab1218 Abcam (Ip), Anti-MDM2 M7815 Sigma, Anti-VHL 68547 Cell Signaling technology.

### Cell proliferation assay

Motor neuron-neuroblastoma hybrid MN-1 cells were seeded on 24-well plate and transfected at 70-80% confluence using Lipofectamine 2000 following manufacturer protocols (Invitrogen). After 48h, 50 μl of MTT (Thiazolyl Blue Tetrazolium Bromide, codice M5655 Sigma-5 mg/ml in PBS1X) was added to each well and the plate incubated for 10’ at 37 °C. Medium was eliminated and DMSO was added to solubilize the water-insoluble MTT formazan crystals. The absorbance was recorded at 570 nm with spectrophotometer Nanodrop2000 (Thermo Fisher).

## Accession numbers

UniProt accession numbers Q00987 and P40337 for the human MDM2 and pVHL proteins, respectively.

## Acknowledgements

This work was supported by Associazione Italiana per la Ricerca sul Cancro (AIRC) grant IG 2015 ID. 17753 and IG 2019 ID. 23825 to ST. The funders had no role in study design, data collection and analysis, decision to publish, or preparation of the manuscript.

## Author contributions

GM, RL, GS, MP, ST conceived the experiments. AF, GM, RB and DP performed the experiments. AF, GM, GS, RL, analyzed the data. GM, RL, GS, MP and ST wrote the manuscript.

## Additional information

### Data Availability

The datasets generated during and/or analyzed during the current study are available from the corresponding author on reasonable request.

