## Supplementary Files for "The E3 ubiquitin-protein ligase MDM2 is a novel interactor of the von Hippel-Lindau tumor suppressor"

### Supplementary material

#### Supplementary Table 1

| Plasmid Name | Primer Name | n° | sequence (5'-3') | Protein expressed |
|---|---|---|---|---|
| pGAD – pVHL30 | VHL30 For | 1 | ggaggccagtgaattcATGCCCCGGAGGGCGGAGAA | Gal4AD-pVHL30 (1-213) |
| | VHL30Rev | 2 | cacccgggtggaattgTCAATCTCCCATCCGTTGAT | |
| pGAD – pVHL19 | VHL19 For | 3 | ggaggccagtgaattcATGGAGGCCGGGCGGCCGCG | Gal4AD-pVHL19 (54-213) |
| | VHL30Rev | 2 | | |
| pGAD – pVHL β | VHL19 For | 3 | | Gal4AD-pVHLβ (54-157) |
| | VHL-β Rev | 4 | cacccgggtggaattgCTAAGTATACACTGGCAGTG TGATATTGGC | |
| pGAD – pVHL α | VHL-α For | 5 | ggaggccagtgaattcCTGAAAGAGCGATGCCTCC | Gal4AD-pVHLα (158-213) |
| | VHL-α Rev | 6 | cacccgggtggaattgTCAATCTCCCATCCGTTGATG TGC | |
| pGAD – pVHL-Nt | VHL30 For | 1 | | Gal4AD-pVHL-Nt (1-53) |
| | VHL-Nt Rev | 7 | cacccgggtggaattgTCACTCCTCCTCGGCGCCCA | |
| pGAD – pVHL (1-157) | VHL30 For | 1 | | Gal4AD-pVHL (1-157) |
| | VHL157Rev | 8 | cacccgggtggaattg TTAAGTATACACTGGCAGTGTGATATTGG | |
| pGBK – MDM2 (1-491) | MDM2 For | 9 | catggaggccgaattcATGGTGAGGAGCAGGCAAAT GTGCAATACC | Gal4BD-MDM2 (1-491) |
| | MDM2 Rev | 10 | ggatccccgggaattgttaGGGGAAATAAGTTAGCAC AATCATTTGAATTGG | |
| pGBK – MDM2 (1-150) | MDM2 For | 9 | | Gal4BD-MDM2 (1-150) |
| | MDM2 (1-150) Rev | 11 | ggatccccgggaattgTTATGAAGATGAAGGTTTCTC TTCCTGAAGC | |
| pGBK – MDM2 (151-350) | MDM2 (151-350) For | 12 | catggaggccgaattcCATTTGGTTTCTAGACCATCT ACCTC | Gal4BD-MDM2 (151-350) |
| | MDM2 (151-350) Rev | 13 | ggatccccgggaattgTTAGTTTTCCAGTTTGGCTTT CTCAGAGATTTCC | |
| pGBK– | MDM2 (351-491) For | 14 | catggaggccgaattcTCAACACAAGCTGAAGAGGG CTTTGATGTTCC | Gal4BD-MDM2 (351-491) |

| | | | | |
|---|---|---|---|---|
| MDM2 (351-491) | MDM2 Rev | 10 | | |
| pGBK – MDM2 (434-491) | MDM2 (434-491) For | 15 | catggaggccgaattcGCCATTGAACCTTGTGTGATTTGTCAAGG | Gal4BD-MDM2 (434-491) |
| | MDM2 Rev | 10 | | |
| pGBK – MDM2 (351-433) | MDM2 (351-491) For | 14 | | Gal4BD-MDM2 (351-433) |
| | MDM2 (351-433) Rev | 15 | ggatccccgggaattgTTAATTAAGGGGCAAACTAGATTCCACACTC | |
| pGBK – MDM2 (351-452) | MDM2 (351-491) For | 14 | | Gal4BD-MDM2 (351-452) |
| | MDM2 (351-452) Rev | 16 | ggatccccgggaattgTTAATGGACAATGCAACCATTTTTAGGTCG | |
| pGBK – MDM2 (329-433) | MDM2 (329-433) For | 17 | catggaggccgaattcTGGGCCCTTCGTGAGAATTGGCTTCC | Gal4BD-MDM2 (329-433) |
| | MDM2 (351-433) Rev | 15 | | |
| pGBK – MDM2 (329-452) | MDM2 (329-433) For | 17 | | Gal4BD-MDM2 (329-452) |
| | MDM2 (351-452) Rev | 16 | | |
| pCDNA 3.1 flag-MDM2 | | | Purchased by Genescript cat n: OHu28568 | Flag-MDM2 (1-491) |
| pCDNA 3.1 flag-VHL30 | | | Purchased by Genescript cat n: OHu23297 | Flag-VHL30 (1-213) |
| pCDNA 3.1 HA-VHL30 | HA-VHL For | 18 | taccgagctcggatcATGGAGTACCCATACGACGTACCAGATTACG | HA-VHL30 (1-213) |
| | HA-VHL Rev | 19 | gatatctgcagaattTCAATCTCCCATCCGTTGATGTGCAATGCGC | |
| | HA-VHL For | 18 | | |

| | | | | |
|---|---|---|---|---|
| pCDNA 3.1 HA-VHL19 | HA-VHL Rev | 19 | | HA-VHL19 (54-213) |

**Table S1. Oligonucletides used in this study.**

List of primers used for vector cloning and construction.

**Supplementary Figure 1**

**A**

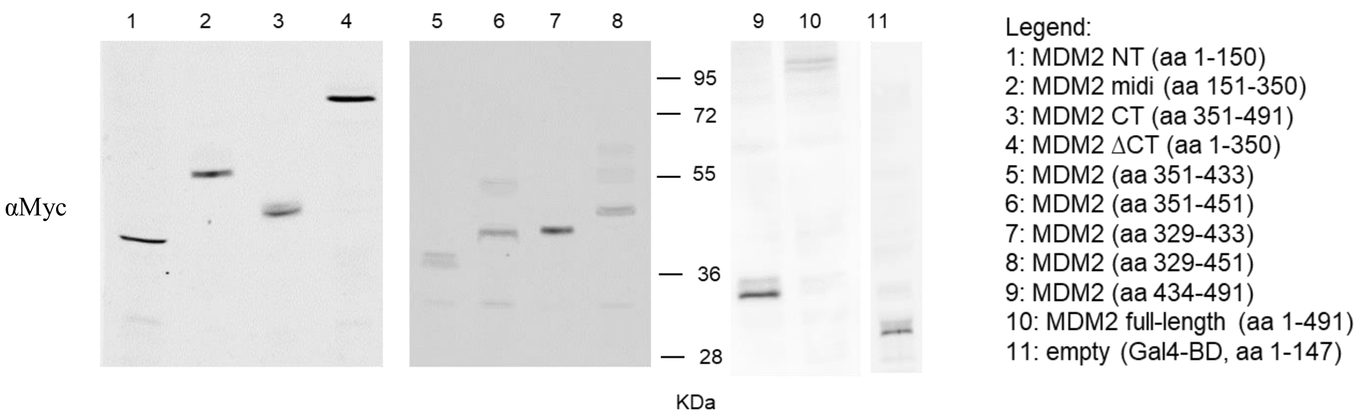


**B**

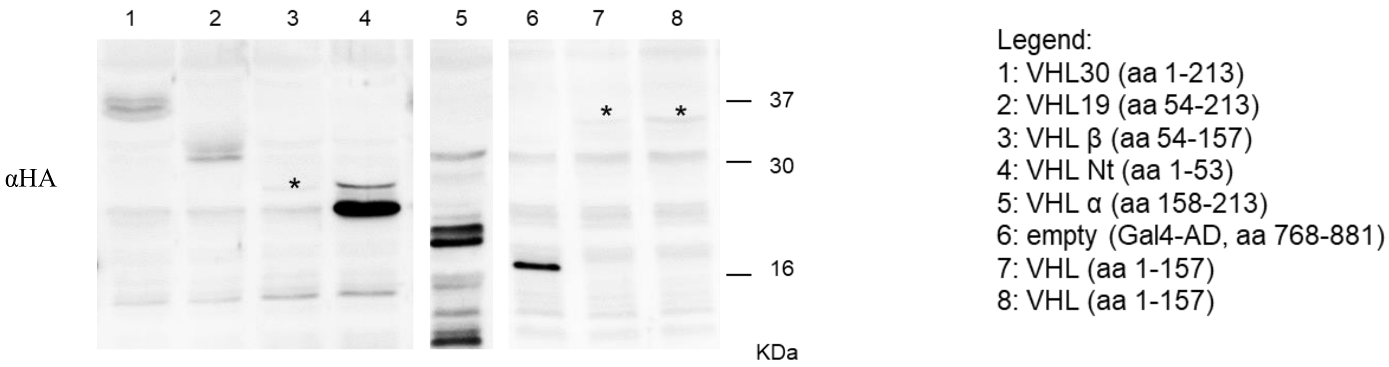


**Figure S1. Expression of Gal4-fusion proteins in yeast.**

A) Western blotting (WB) analysis of total protein extracts of yeast strains expressing the Myc-tagged MDM2 full length and fragments fused to Gal4 DNA binding domain (Gal4-BD). MDM2 has a predicted MW of 55 kDa, yet in WB it runs around 90 kDa, as previously reported (Erhardt. P.,et al. 1997 JBC 272:15049-15052). Gal4-BD + linker have a calculated MW of 21 kDa and runs around 28 KDa. As expected, full length MDM2 run around 110 kDa. Similarly, all fragments in WB have a MW higher than MW calculated.

B) Western blotting (WB) analysis of total protein extracts of yeast strains expressing HA-tagged pVHL full length and fragments fused to Gal4 DNA activation domain (Gal4-AD).

Myc-tagged and HA-tagged proteins were recognized with specific anti-Myc and anti-HA antibodies. Shown is one experiment representative of 3.

**Supplementary Figure 2**

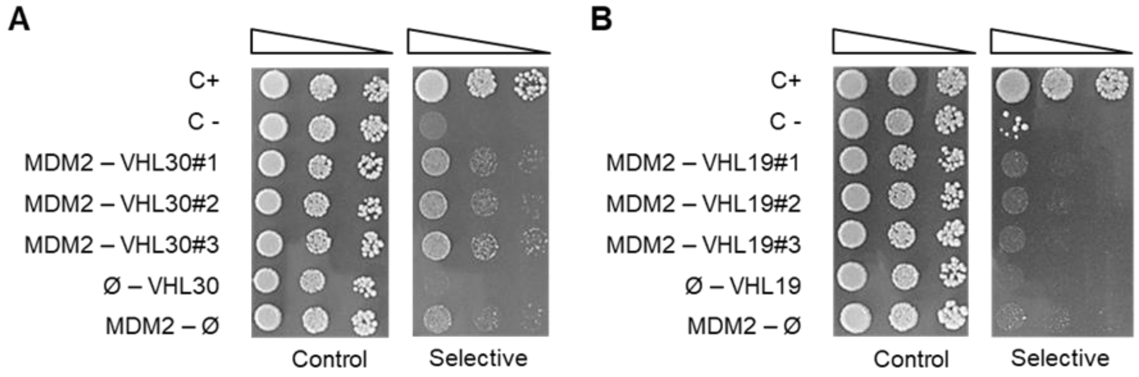


**Figure S2. pVHL30 forms a complex with MDM2.**

A-B) Y2H assays between full length pVHL30 and MDM2 (A), and full length pVHL19 and MDM2 (B) showed that pVHL30 specifically forms a complex with MDM2. (C+) positive control, (C) negative control, 3 clones co-expressing MDM2 (bait) and pVHL30 (prey). Expression of pVHL30 and MDM2 alone did not result in yeast growth in non-permissive medium.

Shown is one experiment representative of 3.

**Supplementary Figure 3**

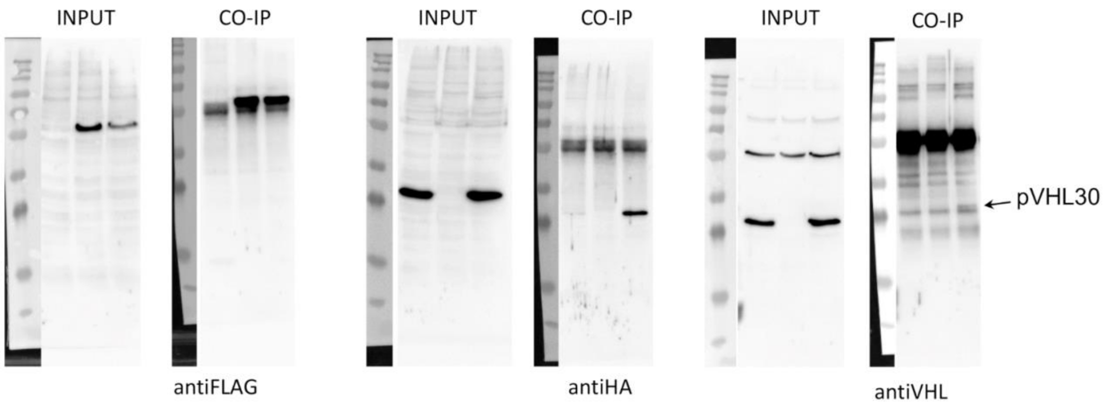


**Figure S3. Uncropped full-length pictures of Western blotting membranes of Figure 2 panel C in the main text**.

**Supplementary Figure 4**

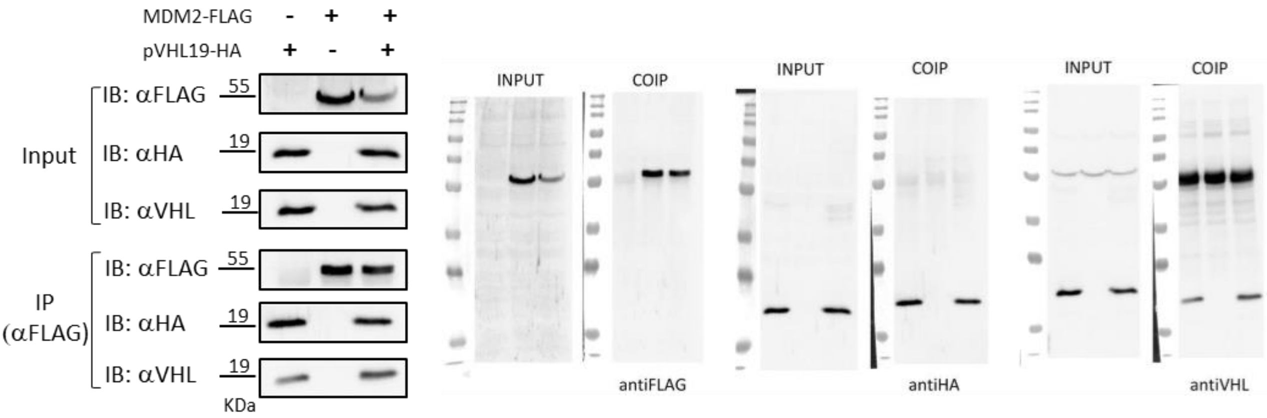


**Figure S4. Immunoprecipitation analysis of MDM2 and pVHL19 interaction in HEK293T cells.** Immunoprecipitation and Western blotting analysis of HEK293T cells expressing Flag-MDM2 and HA-pVHL19. MDM2 was pulled down with an anti-Flag antibody and pVHL19 was analyzed with an anti-HA and pVHL antibodies. The anti-Flag antibody pulled down pVHL19 both in absence and presence of Flag-MDM2, thereby precluding the possibility to test the direct interaction between MDM2 and pVHL19 under these experimental conditions. Shown are cropped and uncropped images from at least 3 independent experiments.

**Supplementary Figure 5**

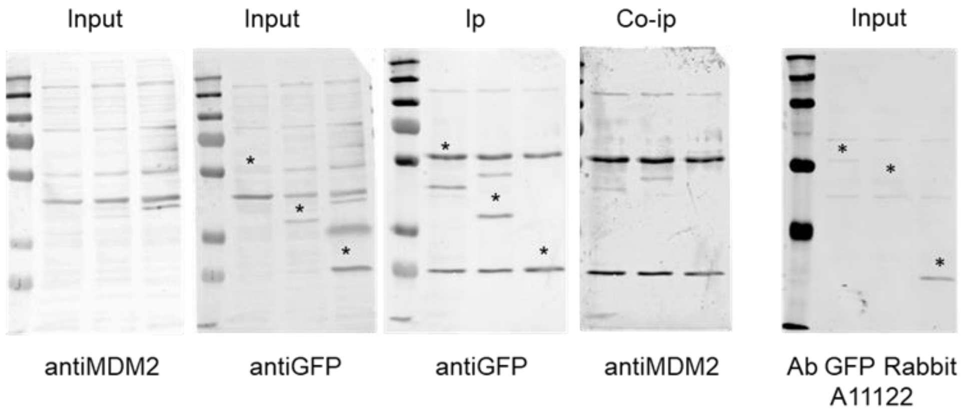


**Figure S5. Uncropped full-length pictures of Western blotting membranes of Figure 2 panel D in the main text**.

Uncropped full-length images of Western blotting membranes of figure 2 panel D in the main text. As additional control inputs were also analyzed using the anti-GFP antibody (A11122, Thermo Fisher Scientific).

**Supplementary Figure 6**

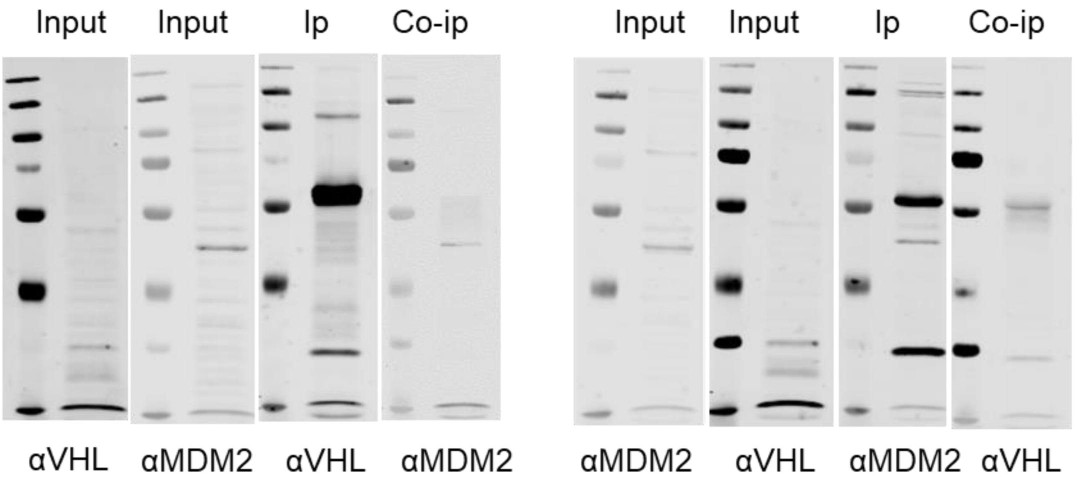


**Figure S6. Uncropped full-length pictures of Western blotting membranes of Figure 2 panel E-F in the main text**.

**Supplementary Figure 7**

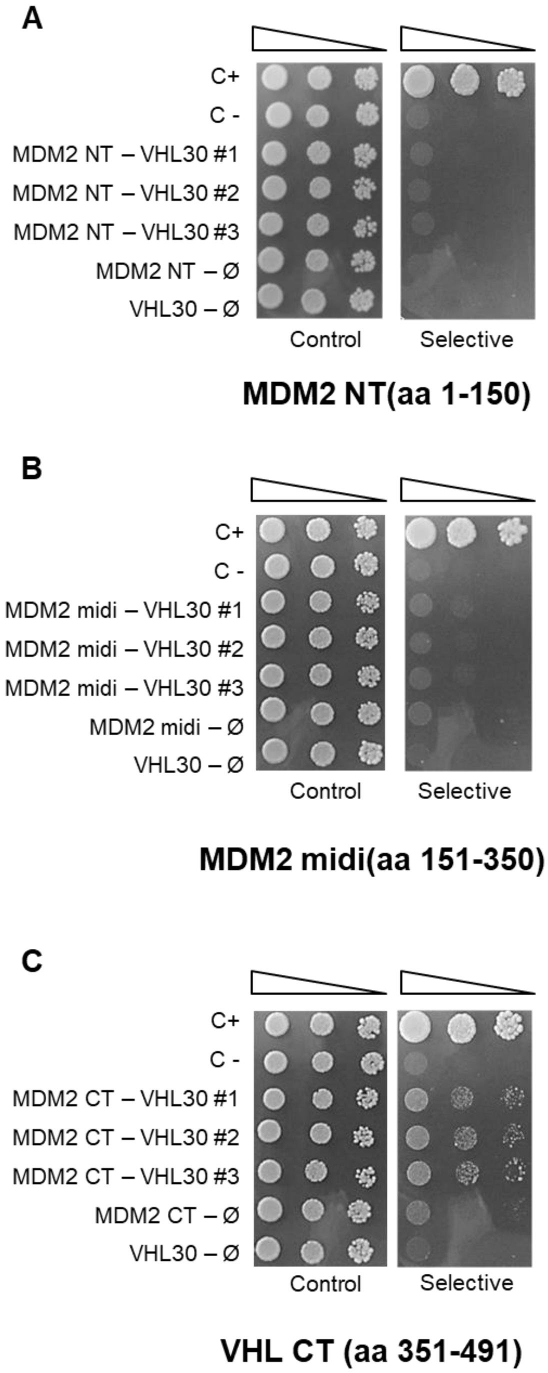


**Figure S7. Uncropped images of figure 3 panel B in the main text. Shown is one experiment representative of 3.**

A: MDM2 NT (aa 1-150); B: MDM2 middle (aa 151-350); C: MDM2 CT (aa 351-491).

**Supplementary Figure 8**

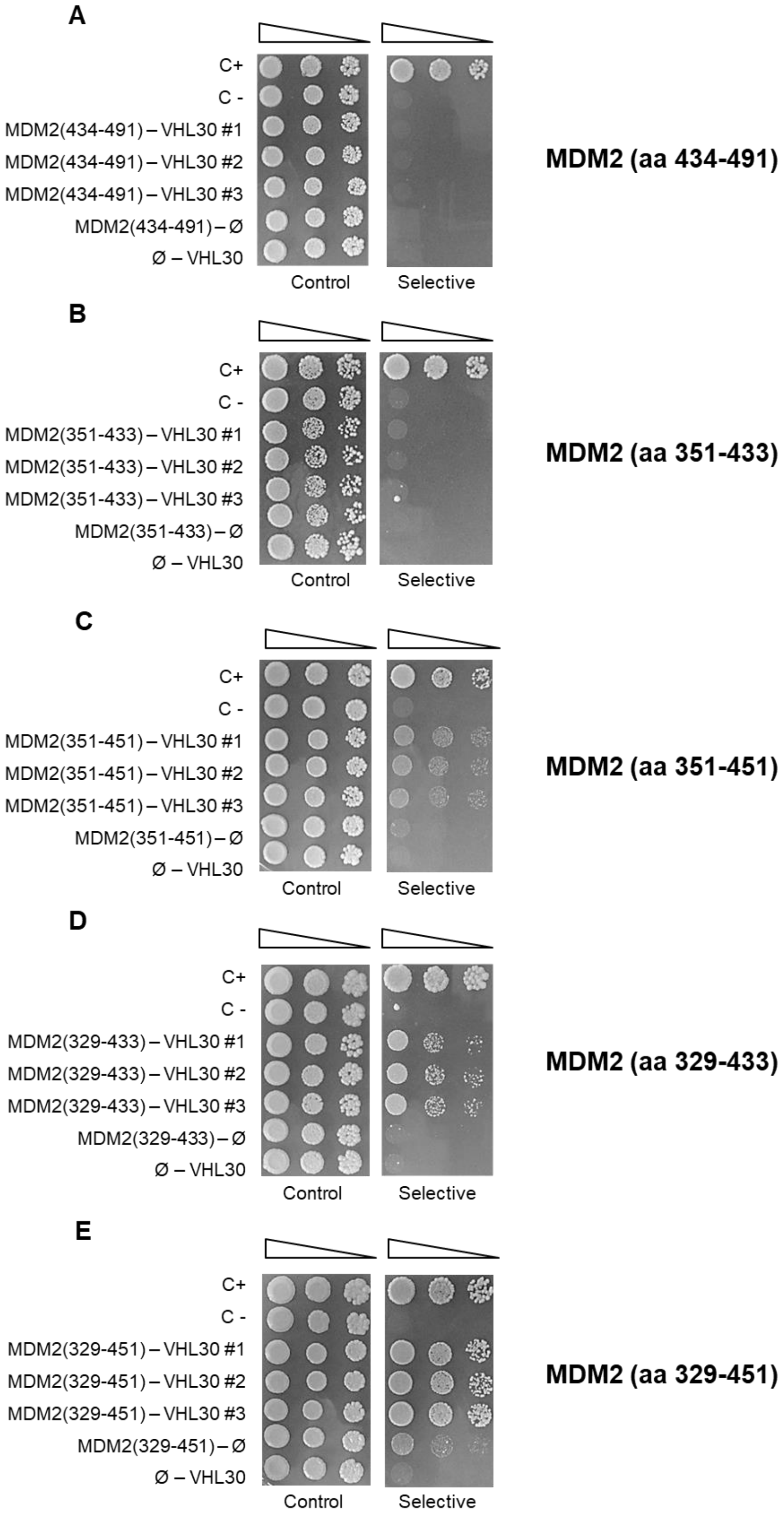


**Figure S8. Uncropped images of figure 3 panel E in the main text. Shown is one experiment representative of 3.**

A: MDM2 (aa 434-491); B: MDM2 (aa 351-433); C: MDM2 (aa 351-451); D: MDM2 (aa329-433) and E: MDM2 (aa 329-451).

**Supplementary Figure 9**

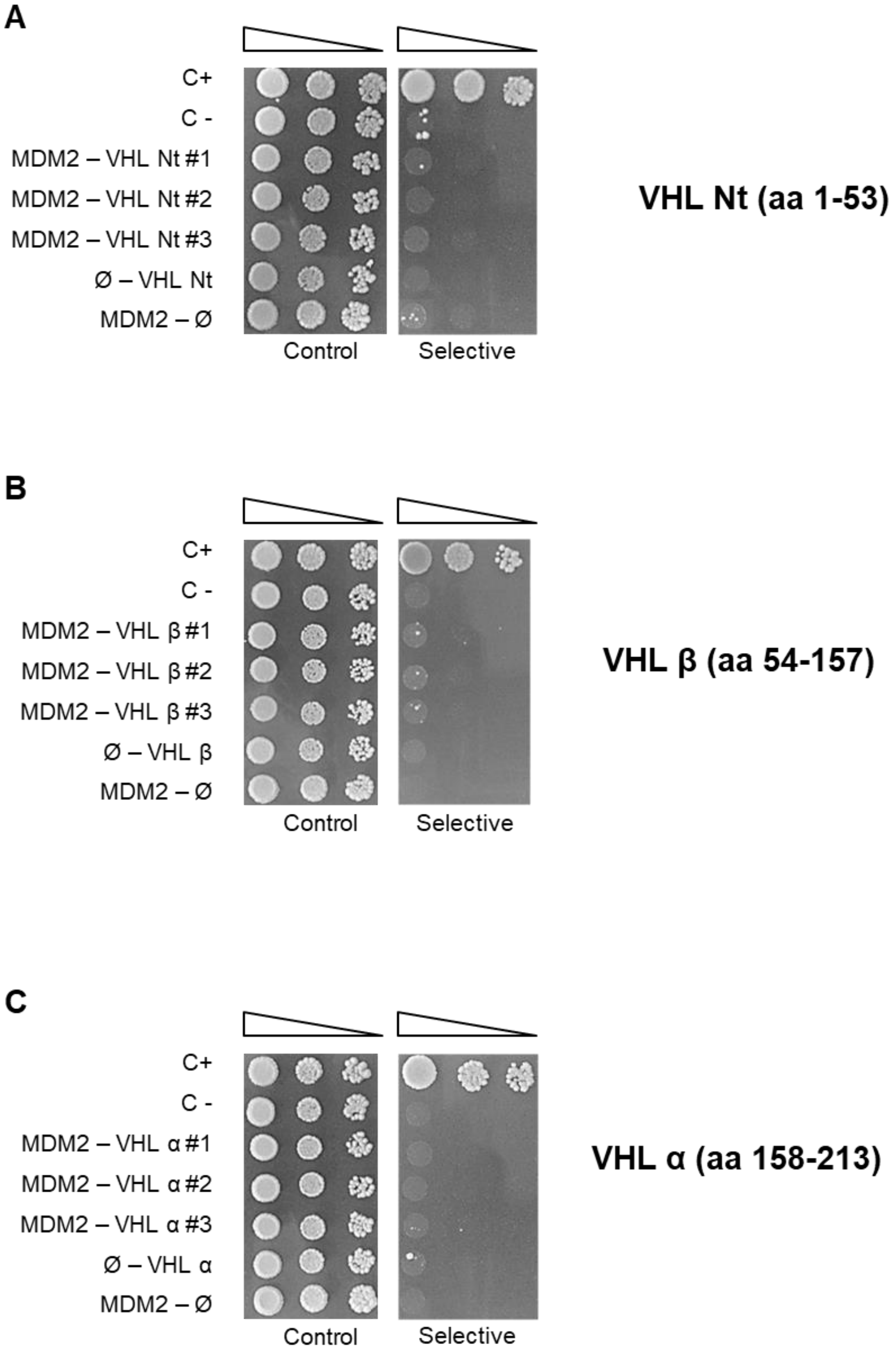


**Figure S9. The N-terminus and β- and α-regions of pVHL30 do not associate with MDM2.** A-C) Y2H assays analysis of interaction between full-length MDM2 and (A) pVHL Nt (1-53), (B) pVHL β domain (aa 54-157), and (C) pVHL α domain (aa 158-213). Shown is one experiment representative of 3.

**Supplementary Figure 10**

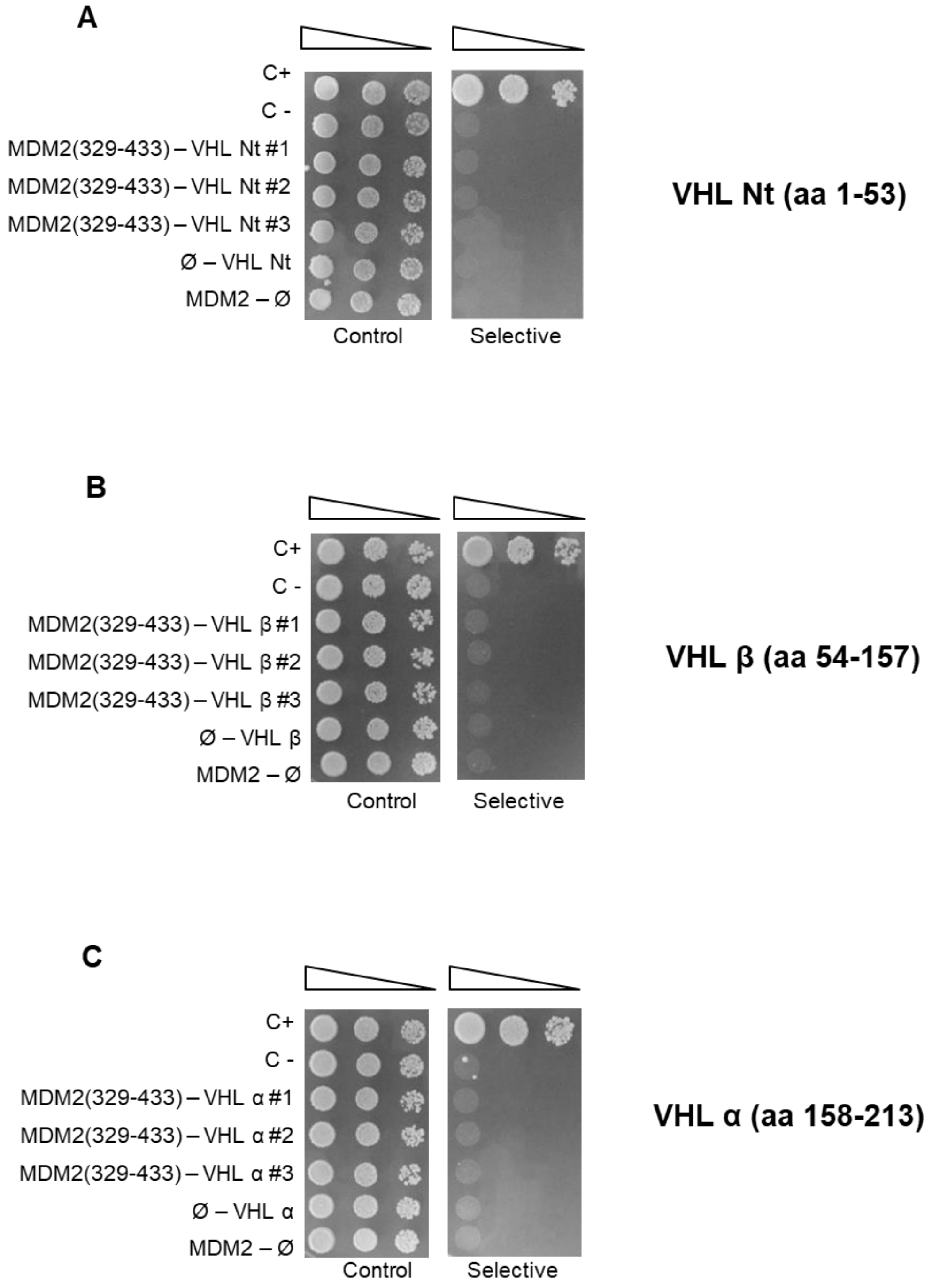


**Figure S10. Uncropped images of figure 4 panel B in the main text. Shown is one experiment representative of 3.**

A: VHL Nt (aa 1-53); B: VHL β domain (aa 54-157); C: VHL α domain (aa 158-213).

**Supplementary Figure 11**

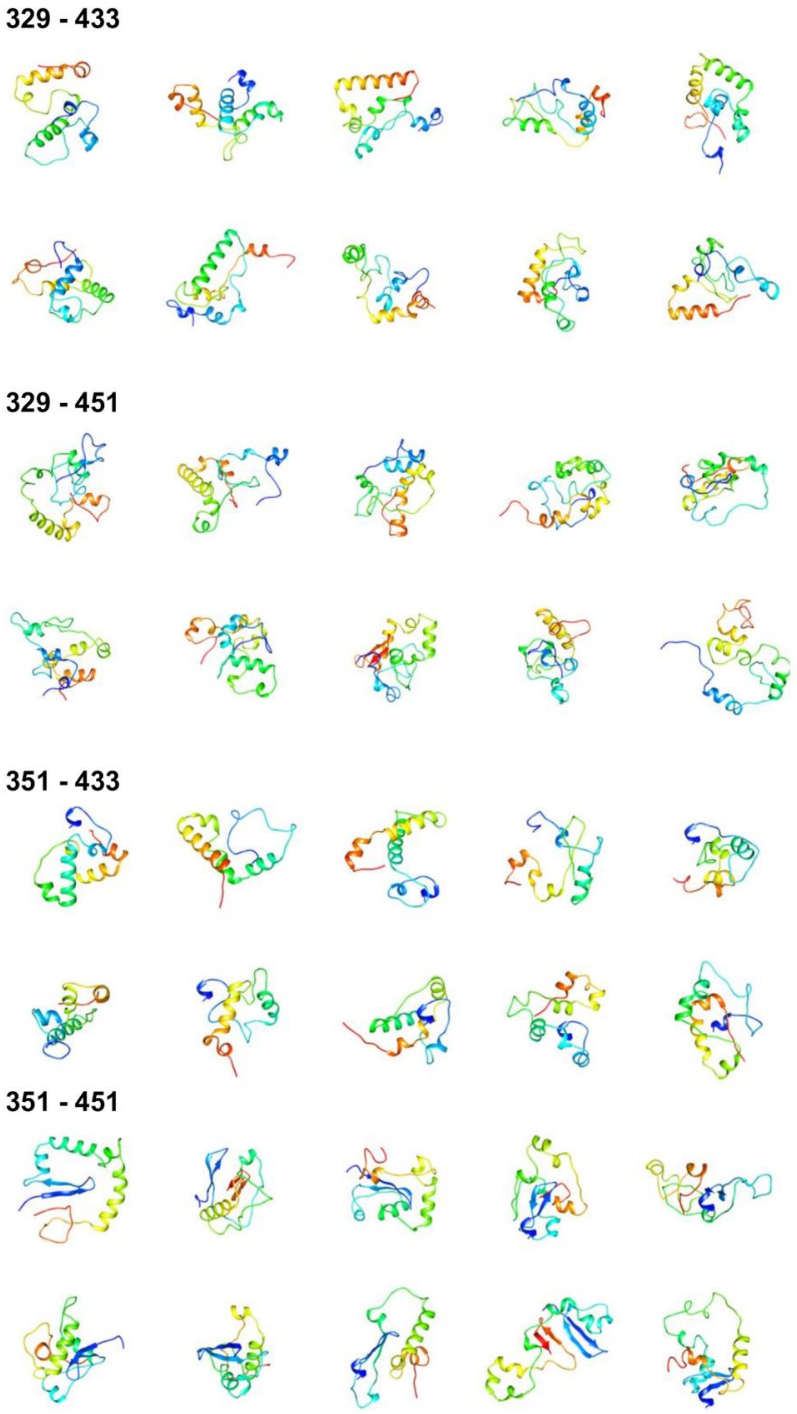


**Figure S11. Overview of predicted 3D structure of each MDM2 fragment.**

The best ten 3D structures calculated for the MDM2 fragments used during yeast two hybrid experiments. Structures are presented as cartoon and colored blue (N-terminal) to red (C-terminal).

**Supplementary Figure 12**

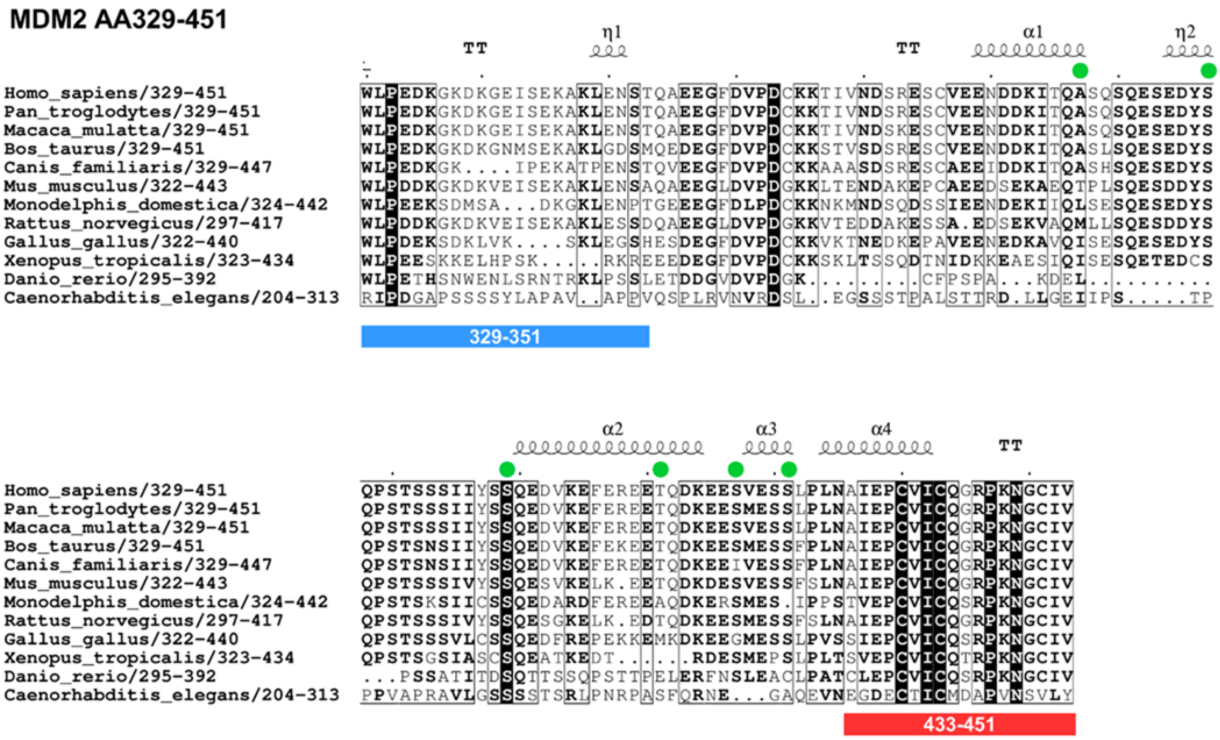


**Figure S12. Multiple sequence alignment of MDM2 329-451 fragment.**

Colored boxes represent the position of the two flanking regions presumed to drive MDM2/pVHL30 association. Green sphere highlight position of phosphorylation sites.

**Supplementary Figure 13**

αMyc

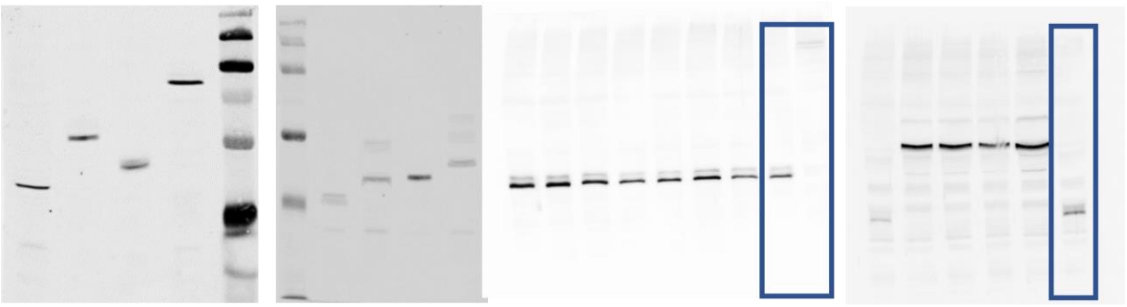

αHA

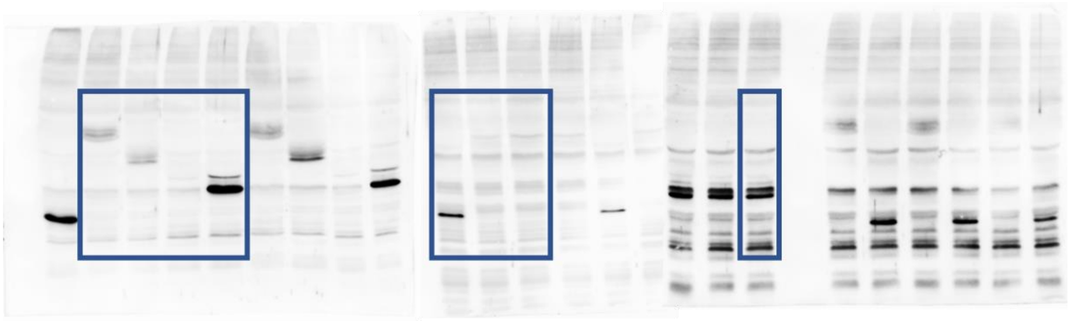

**Figure S13 Western blot**

Uncropped full-length pictures of western blot membranes referred to figure S1.
